# Physical performance and telomere length in older adults

**DOI:** 10.1101/2023.10.30.564820

**Authors:** José Darío Martínez-Ezquerro, Mauricio Ortiz-Ramírez, Paola García-delaTorre, Vanessa González-Covarrubias, Sergio Sánchez-García

## Abstract

**Background:** The fast-paced aging demographic prompts studying risk factors and markers that could predict healthy aging. Telomere length shows promise for assessing a broad spectrum of aging-related phenotypes.

**Aim of the study:** To assess the association between physical performance and telomere length in Mexican older adults.

**Methods:** In this observational cross-sectional study, we recruited 323 older adults affiliated with the Instituto Mexicano del Seguro Social (IMSS) and members of the “Cohort of Obesity, Sarcopenia, and Frailty of Older Mexican Adults’’ (COSFOMA). To assess physical performance, we used the Short Physical Performance Battery (SPPB) and stratified individuals into two groups according to their SPPB score into low ≤7 (L-SPPB) and high >7 (H-SPPB). Absolute telomere length (aTL) was determined by using qPCR. Next, we classified individuals according to their aTL into short ≤4.22 kb (S-TL) and long >4.22 kb (L-TL). For both SPPB and aTL categories, we calculated the mean and adjusted mean by sex, age, marital status, education, remunerated work, smoking, drinking, cognitive decline, depression, and polypharmacy with 95% CI, including the lower and upper CI (_LCI_mean_UCI_). We estimated the effect size between physical performance and telomere length with Cohen’s d for unequal group sizes. Also, we calculated the odds ratio with 95% confidence intervals, including the lower and upper CI (_LCI_OR_UCI_) for physical performance according to telomere length categories.

**Results:** Participants in the low physical performance category had significantly shorter telomeres (_4.1_4.4_4.7_ mean and _3.5_4.0_4.5_ kb adjusted mean, p<0.001), in opposition to the high physical performance category (score >7) with longer telomeres (_5.5_5.7_5.9_ mean and _4.7_5.3_5.8_ kb adjusted mean, p<0.001), with a medium-to-high telomere length effect size (d= 0.762). Finally, the odds of being classified in the low physical activity category increased _2.1_3.6_6.1_ times per kb of telomere (adjOR _1.7_3.3_6.3_, p<0.001) compared to the high physical activity group (p<0.001).

**Conclusion:** Decreased physical functioning is associated with lower telomere length. Absolute telomere length (aTL) as a possible biomarker for differential diagnosis of healthy and unhealthy aging should be explored further.

## Introduction

### Aging & Healthy Aging

At the beginning of 2021, there were over one billion older adults worldwide, which is estimated to double by 2050 [1]. Old age refers to the final stage of life and it is characterized by phenotypic changes resulting from allostasis and other changes that occur throughout life, where the resulting modifications of the adaptive capacity become evident [2]. Aging is characterized by a progressive loss of physical integrity, leading to impaired function and increased vulnerability to death [3]. The heterogeneous aging process among individuals is intricately linked to all biological levels, from molecules to ecosystems, contributing to its complexity [4]. Common characteristics of aging include genomic instability, telomere attrition, epigenetic alterations, loss of proteostasis, disabled macroautophagy, deregulated nutrient sensing, mitochondrial dysfunction, cellular senescence, stem cell exhaustion, altered intercellular communication, chronic inflammation, and dysbiosis [3,5]. Alterations in these hallmarks of aging generate the clinical phenotypes observed in older persons: anemia, a blunted immune response system, cognitive impairment, delirium, depression, frailty, hearing and visual impairment, mobility impairment and falls, osteoporosis, poor nutrition, pressure score, sarcopenia, sleep disorders [6,7]. Since 2015, the World Health Organization (WHO) defines healthy aging as the process of maintaining functional ability to enable well-being in older age. This person-centered approach considers interactions between both life trajectories and dynamic environments to determine functional ability [8].

### Physical performance and SPPB

Physical performance can be defined as the functional ability of individuals to perform certain physical activities that depend on a variety of factors that positively or negatively affect their health; therefore, it is considered an indicator of health status [9].

One well-known instrument to measure physical function is the Short Physical Performance Battery (SPPB), which can be particularly useful to assess lower limb function. This standardized test comprises three assessment domains: standing balance, walking speed, and five times sit-to-stand (5xSTS) tests [10]. The SPPB is commonly used in geriatric evaluations for its ability to predict subsequent health-related outcomes, including falls and disability [11,12]. This instrument assesses standing balance as the ability to stand for up to 10 seconds with feet positioned in three ways (together side-by-side, semi-tandem, and tandem) [13]. Walking speed is measured with the 4-meter gait speed test by a stopwatch [14]; the time required for a person to walk a distance without obstacles, being able to use assistive devices if needed [12], is one of the most widely used assessment tools and most common way to assess physical performance [14]. Finally, performance on the 5xSTS task is measured as the time required to transit from a sitting position to standing five times without using the upper extremities [10,15,16]; this assessment has been previously associated with future falls and disability in older adults [11].

### Telomere length: normal and accelerated attrition

Telomere length may serve as a potential biomarker to assess, predict, and monitor overall health. Telomere shortening is associated with aging-related conditions, including cardiovascular disease [17], diabetes mellitus [18], some types of cancer [19], physical activity [20], and Alzheimer’s disease [21].

Telomeres are non-coding regions of the DNA sequence TTAGGG repeated in tandem [22] with a length of 5-15 kb in humans which are subject to shortening during each cell division, varying with increased age [23], and located at the ends of linear chromosomes [24] in three-dimensional arrangements that confer protection to chromosomes and prevent their recognition as DNA damage, activation of apoptosis, DNA degradation as well as fusion with other neighboring chromosomes [23,25]. Their length can be measured by methods based on binding nucleic acid probes or primers to telomere-specific DNA repeat sequences [26].

Normally, the telomeres in somatic cells shorten at each cell division due to a phenomenon known as the end-replication problem, where small DNA fragments of approximately 50 bp of telomere sequence are lost in each cell division [27]. Telomere attrition is a natural cellular process that occurs over time; however, certain factors can accelerate their shortening, such as chronic stress [28], inflammatory processes and lack of physical activity [29], polyunsaturated fatty acid intake [30], smoking [31], environmental pollutants [32], NO2 [33], and organophosphate insecticides [34]. All these factors can generate oxidative stress at the cellular level, leading to cellular aging [35–37]. Consequently, cells enter a state of senescence, undergo morphological and genetic changes that lead to tissue dysfunction [38], contributing to functional decline and the aging process [39].

### Physical performance and telomere length

Physical activity has been associated with a healthy lifestyle [40], as well as with a decreased risk of suffering from some chronic degenerative diseases such as type 2 diabetes [41], hypertension [42], and cancer [43] in older adults.

A physiological way to explain this decrease in risk could be through telomere length since it is considered a marker of aging and has been associated with a healthy lifestyle and physical activity [44]. When physical activity is performed, an antioxidant and anti-inflammatory effect is generated in the body [45], which can delay telomere shortening [46]. However, the association between telomere length and physical activity remains uncertain due to the contradictory results reported by various studies [47]. While some studies report an association between telomere length and physical activity [48–50], other authors have described no association between physical activity and telomere length [51,52]. Of note, at least one unexpected result from a study in older Brazilian subjects has shown the opposite: better physical capacity associated with lower telomere length [53].

Given these observations, the aim of this study was to evaluate the association between physical performance and telomere length in older Mexican adults and contribute to the understanding and further disambiguation of the contradictory results reported previously.

We hypothesize that the most biologically plausible association would be an accelerated telomere length attrition —that may be indirectly observed as shorter telomere length— in participants with low physical performance compared to those with high physical function.

## Methods

### Participants

In this cross-sectional study, we invited older residents of Mexico City, beneficiaries of Instituto Mexicano del Seguro Social (IMSS), members of the COSFOMA cohort (Cohort of Obesity, Sarcopenia and Frailty of Older Mexican Adults). The COSFOMA cohort comprised 1,252 adults aged 60 years and older, from both Mexico City and Estado de México. This research protocol underwent a comprehensive review and obtained approval from the IMSS National Scientific Research Committee, which includes the Health Research Committee, Health Research Ethics Committee, and Health Research Biosafety Committee. The study was assigned the registry number R-2012-785-067. Participants of the COSFOMA cohort were initially contacted through an informative invitation letter to participate in this study. Their agreement to participate was subsequently confirmed via telephone. Those who agreed to participate provided written informed consent before any research-related procedures were conducted [54].

### Physical performance assessment

To evaluate lower extremity functioning in older adults, we used an objective assessment tool, the Short Physical Performance Battery (SPPB) [55,56] https://www.nia.nih.gov/research/resource/short-physical-performance-battery-sppb, a standardized test composed of three assessment domains: *Standing balance*: assessed as the ability to stand for up to 10 seconds with feet positioned in three ways (together side-by-side, semi-tandem, and tandem) [13] *Walking speed*: measured with the 4-meter gait speed test by a stopwatch [14] *Five times sit-to-stand (5xSTS) tests*: measured as the time required to transit from a sitting position to standing five times without using the upper extremities [10,15,16]. Each task has a score range between 0-4 points, which is used to summarize the physical performance score (SPPB; 0-12 points) [10].

We used SPPB scores to categorize participants into two groups, considering the bottom 20th centile: low (L-SSPB; score ≤7, n=104) and high (H-SPPB; score >7, n=219) physical performance.

### Telomere length

We obtained peripheral blood samples by venipuncture, purified DNA from leukocytes following the salting-out method [57], and stored 10 ng/μL aliquots at −70°C until analysis. To measure absolute telomere length (aTL), we followed O’Callaghan and Fenech method [58] with real-time quantitative polymerase chain reaction using Maxima SYBR Green/ROX qPCR Master Mix 2X (Thermo Scientific, Waltham, MA, USA) reaction (conditions: 10 min at 95°C, followed by 40 cycles of 95°C for 15s, 60°C for 1 min) followed by a melting curve to detect artifact amplification. To assess telomere length in base pairs, we generated a calibration curve with serial dilutions (10^-1^ through to 10^-6^ dilutions) of both a synthesized oligomer standard (STD) of fourteen TTAGGG repeats (84-mer telomere STD) and a housekeeping gene reference of 75 nucleotides (75-mer 36B4 STD; CAGCAAGTGGGAAGGTGTAATCCGTCTCCACAGACAAGGCCAGG ACTCGTTTGTACCCGTTGATGATAGAATGGG) [59,60] (Table 1). Then, we calculated both standard curves and extrapolated by linear regression the Ct value of each sample to its corresponding telomere 84-mer repeats and housekeeping gene copy number.

**Table 1.**
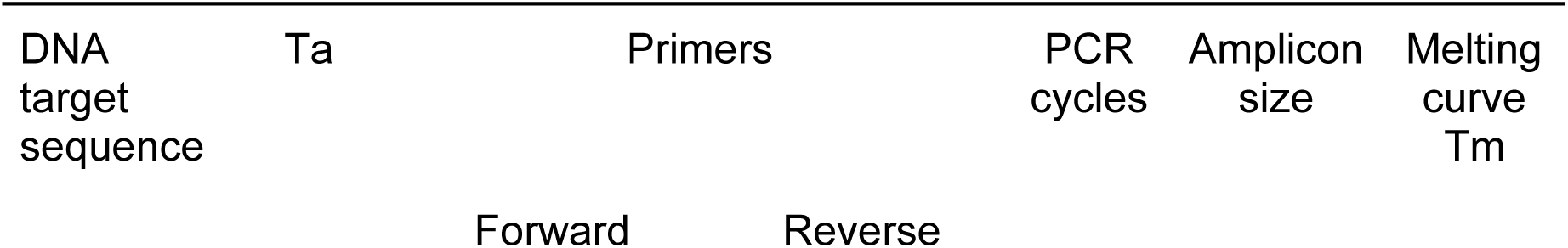
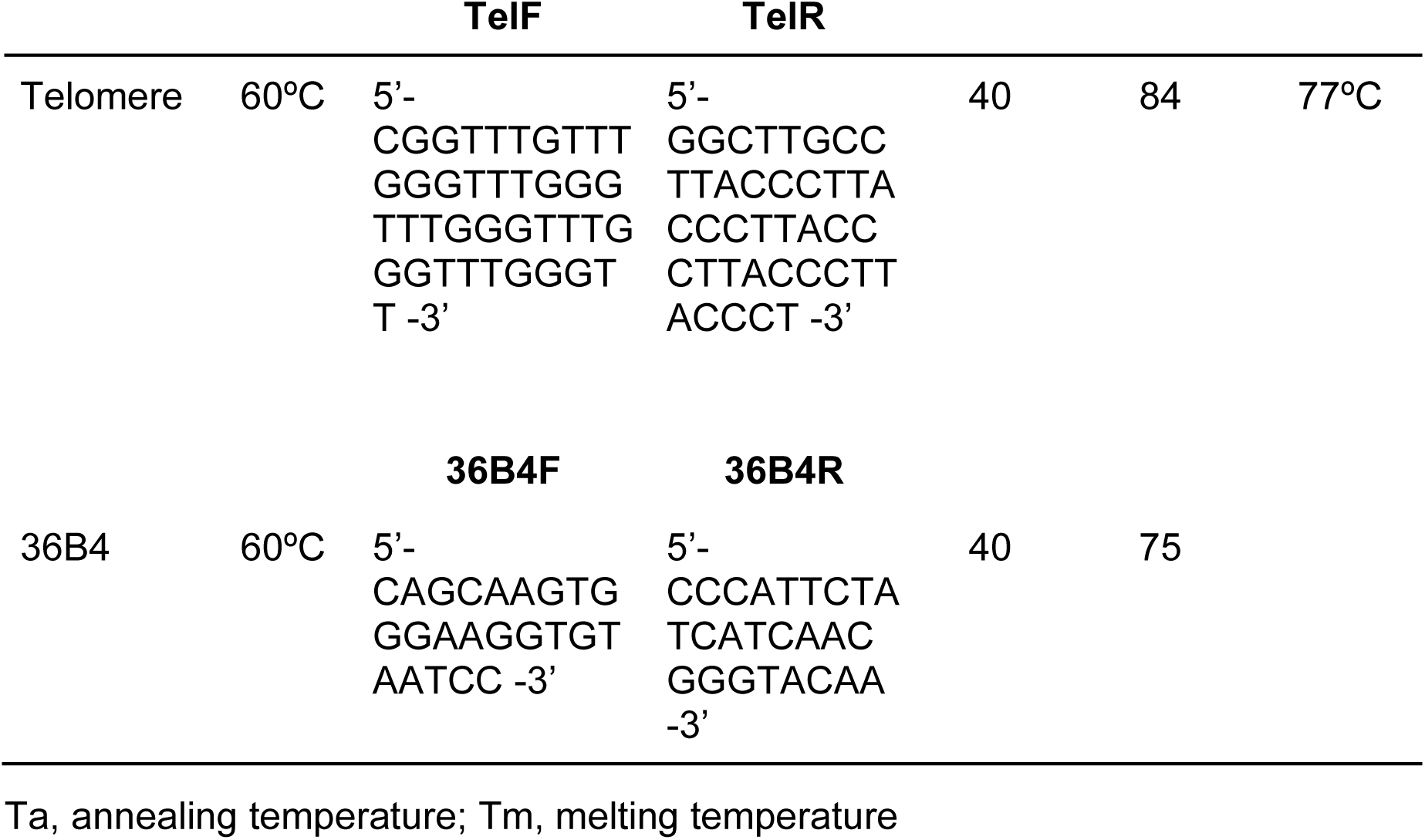
Real-time quantitative polymerase chain reaction (qPCR) and melting curve conditions.

### Statistical analysis

Descriptive analysis of the participant’s characteristics included qualitative variables presented as absolute (n) and relative (%) frequencies and quantitative variables as the arithmetic mean and standard deviation (SD). We obtained telomere length values in base pair units for each sample by extrapolating their Ct value from qPCR regression equations.

We compared differences between telomere length (kb pairs) and physical performance (SPPB scores) according to the general characteristics of participants (Student’s t-test for independent samples for two groups or ANOVA for three groups).

To assess physical performance according to their telomere length, we stratified participants into two SPPB groups: low and high physical performance (L-SPPB score ≤7 and H-SPPB score >7, respectively; bottom 20th centile) and calculated the mean and adjusted mean by sex, age, marital status, education, remunerated work, smoking, drinking, cognitive decline, depression, and polypharmacy with 95% CI, including the lower and upper CI (LCI and UCI, respectively: _LCI_mean_UCI_).

We grouped older adults into short and long telomere length groups (≤4.22 kb and >4.22 kb; bottom 25th percentile, respectively). We calculated the mean and adjusted mean by sex, age, marital status, education, remunerated work, smoking, drinking, cognitive decline, depression, and polypharmacy with 95% CI (_LCI_mean_UCI_) for the SPPB and telomere length categories. To measure the difference between the two group means, we estimated the effect size (Cohen’s d for unequal group sizes) between physical performance and telomere length with an independent samples t-test with 95% CI, including lower and upper CI. We calculated the odds ratio with 95% lower and upper confidence intervals (_LCI_OR_UCI_) for physical performance according to telomere length categories.

## Results

Members from the COSFAMM cohort (Cohort of Obesity, Sarcopenia, and Frailty of Older Mexican Adults, COSFOMA) involving 1,252 adults ≥60 years of age from Mexico City and Estado de México (Figure 1, [61]) were invited by phone to participate in this study. A total of 323 (26%) COSFAMM members agreed to participate: 212 women (65.6%) and 111 men (34.4%) (F/M sex ratio: 212/111), with a mean±SD age of 70.1 ± 7.2 years. Around half of older participants were between 60-69 years of age (52.9%), with normal weight (62.2%), married (53.6%), with seven or more years of education (50.5%), without reported comorbidities (55.1%) or cognitive decline (53.3%), but under polypharmacy (50.8%). Only 28.5% had paid employment, 10.8% lived alone, 7.1% smoked, 18.9% consumed alcohol, and 38.4% were depressed. The general characteristics of the participants by physical performance and telomere length groups are summarized in Table 2.

**Figure 1.**
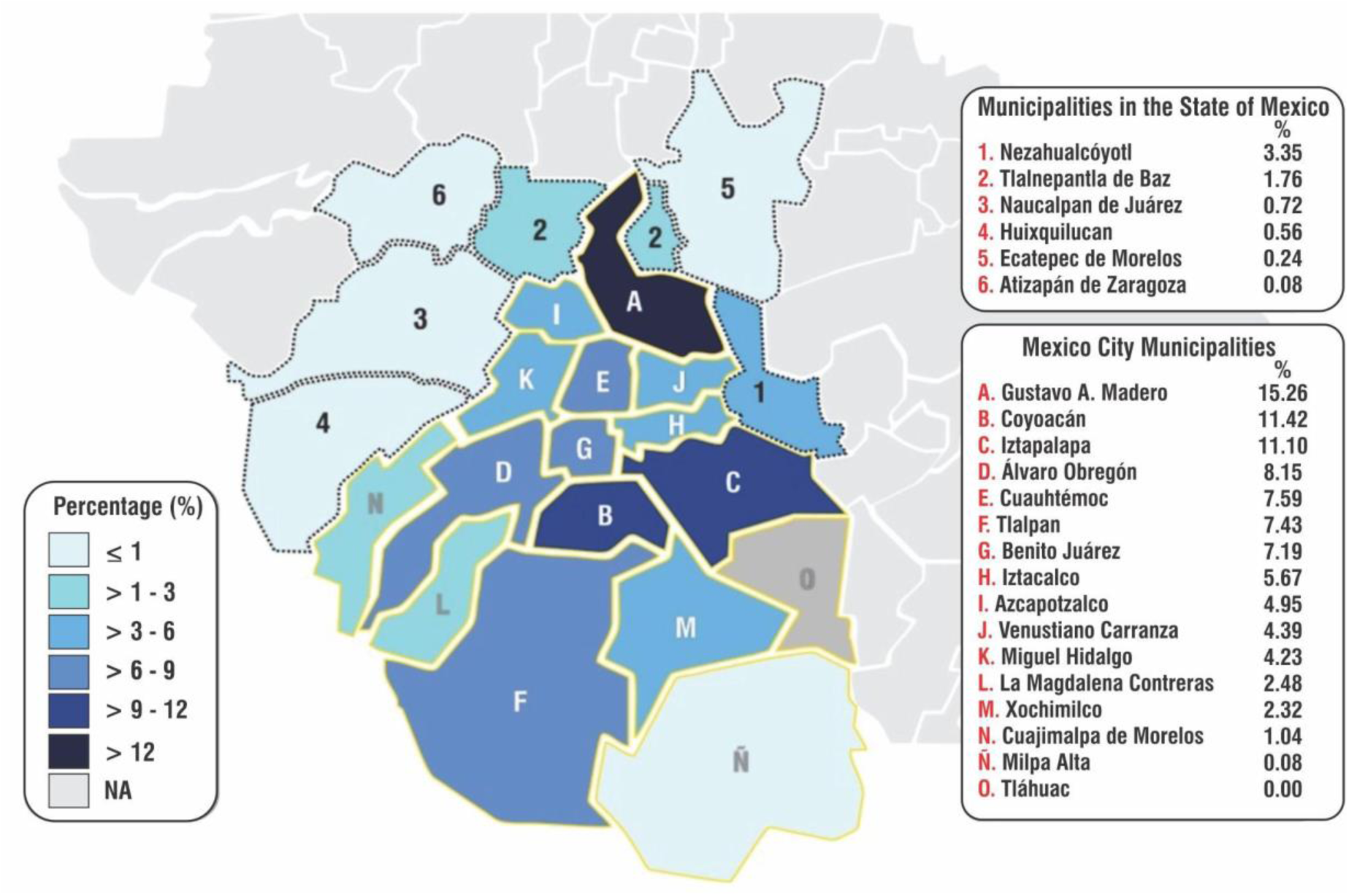
Geographical distribution of the COSFAMM cohort participants. The map shows the percentage geographic distribution of the cohort participants (n= 1,252; ≥60 years of age) by municipalities from two Mexican states, Estado de México (gray divisions and numbers) and Mexico City (CDMX; yellow division and letters) [61].

**Table 2.**
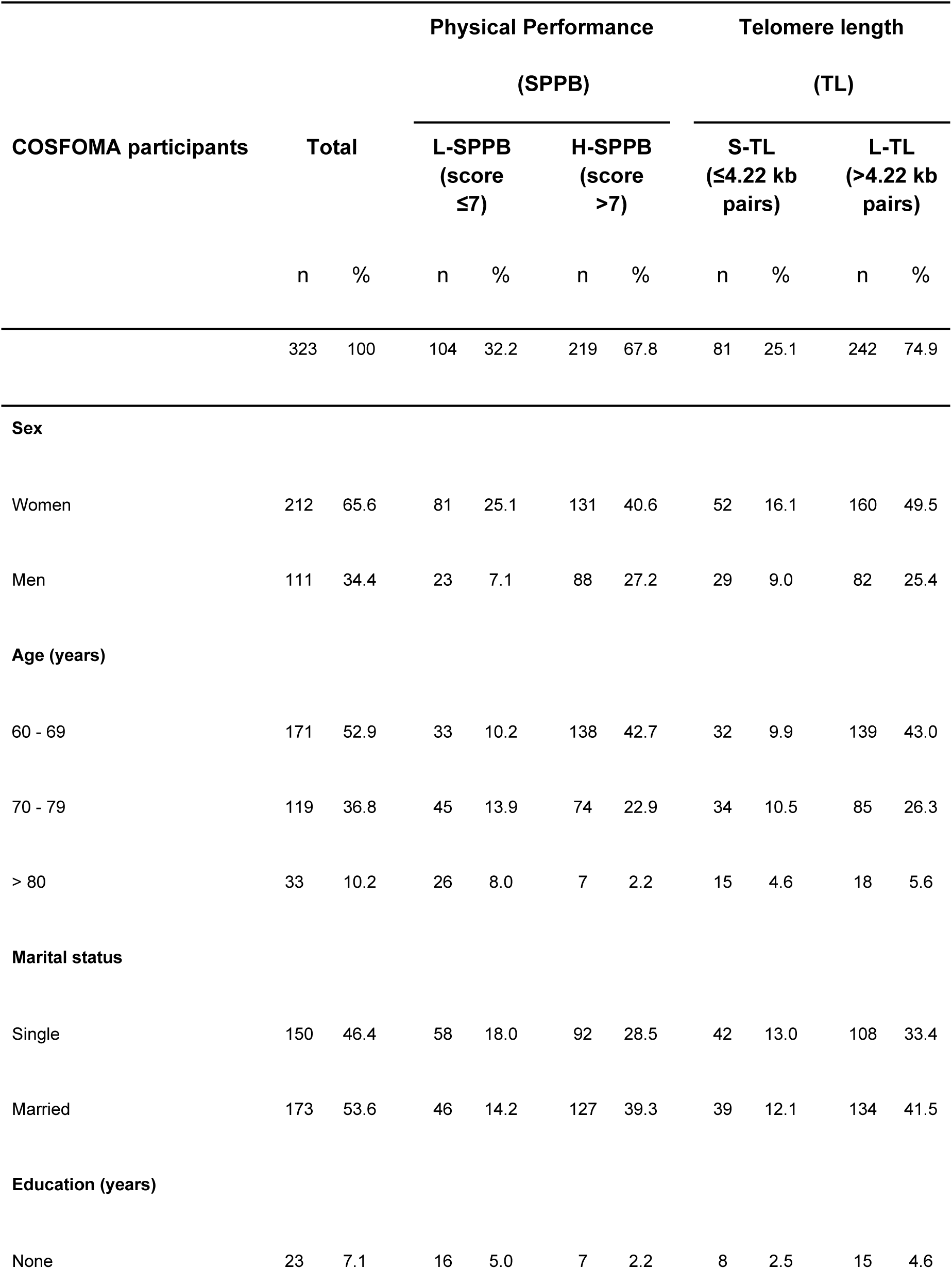
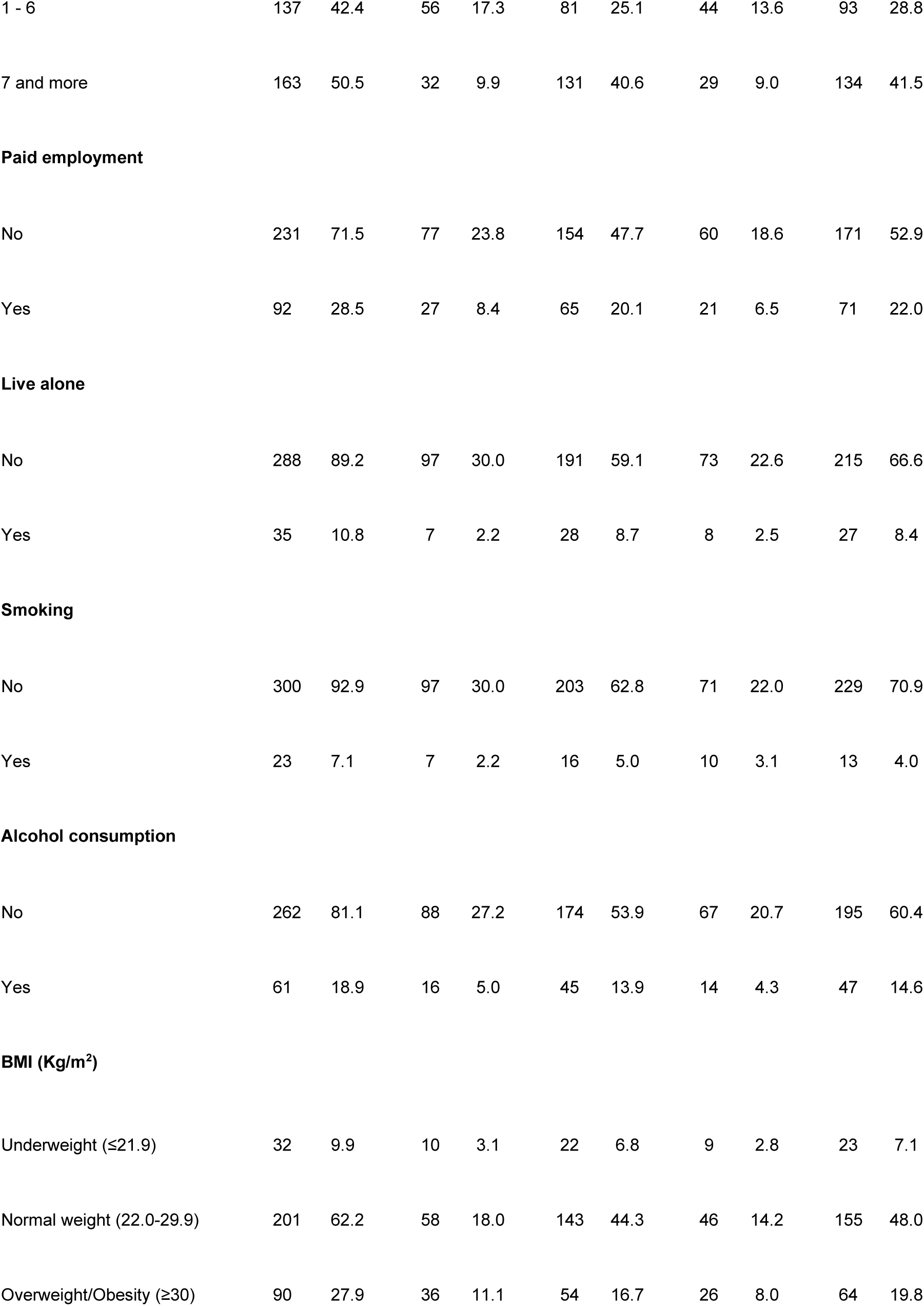
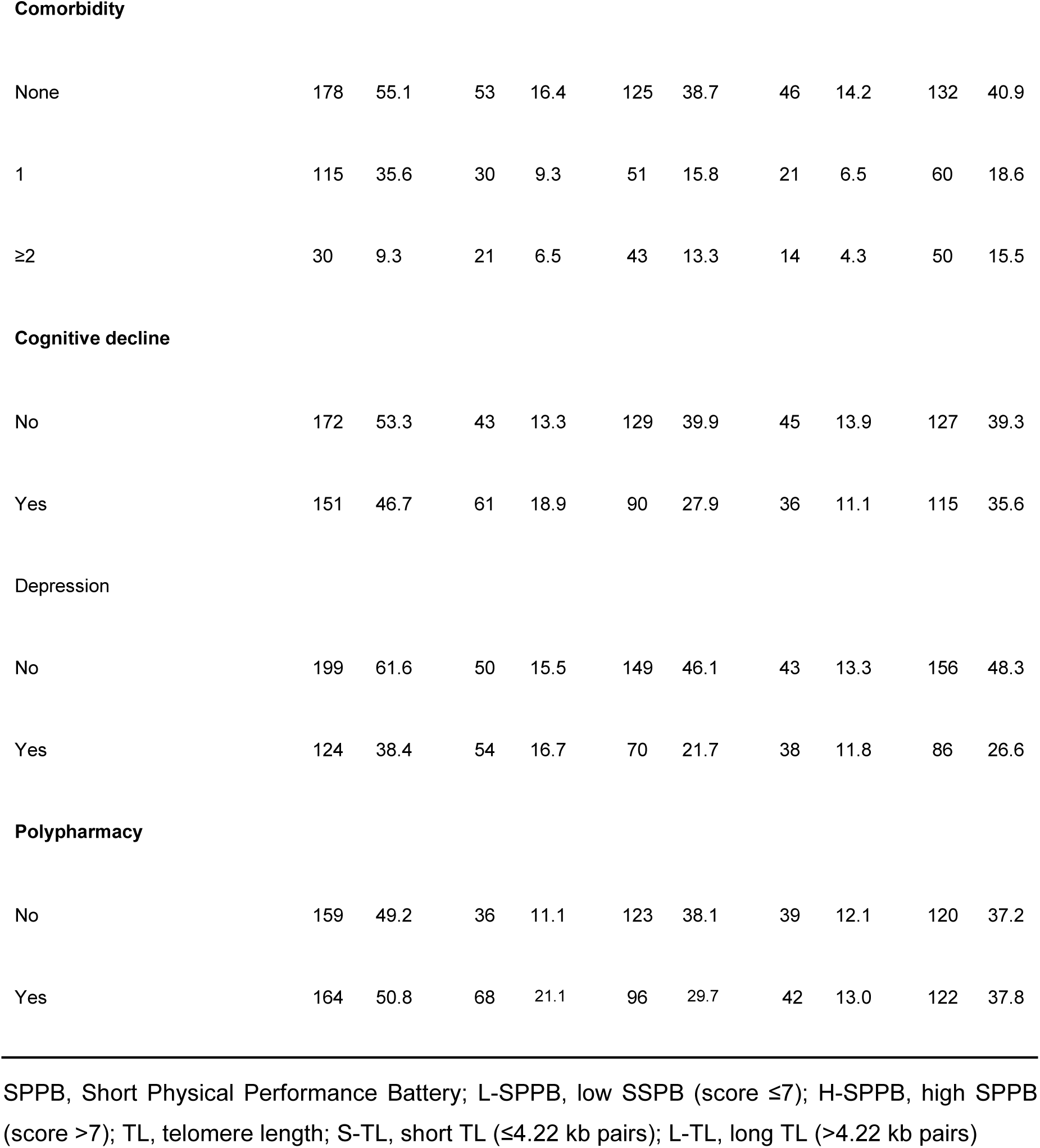
General characteristics of COSFOMA participants (n=323).Table 3. Physical performance categories by short and long telomere length.

| COSFOMA participants | Total |  | Physical Performance |  |  |  | Telomere length |  |  |  |
| --- | --- | --- | --- | --- | --- | --- | --- | --- | --- | --- |
|  |  |  | (SPPB) |  |  |  | (TL) |  |  |  |
|  |  |  | L-SPPB<br>(score ≤7) |  | H-SPPB<br>(score >7) |  | S-TL<br>(≤4.22 kb pairs) |  | L-TL<br>(>4.22 kb pairs) |  |
|  | n | % | n | % | n | % | n | % | n | % |
|  | 323 | 100 | 104 | 32.2 | 219 | 67.8 | 81 | 25.1 | 242 | 74.9 |
| **Sex** |  |  |  |  |  |  |  |  |  |  |
| Women | 212 | 65.6 | 81 | 25.1 | 131 | 40.6 | 52 | 16.1 | 160 | 49.5 |
| Men | 111 | 34.4 | 23 | 7.1 | 88 | 27.2 | 29 | 9.0 | 82 | 25.4 |
| **Age (years)** |  |  |  |  |  |  |  |  |  |  |
| 60 - 69 | 171 | 52.9 | 33 | 10.2 | 138 | 42.7 | 32 | 9.9 | 139 | 43.0 |
| 70 - 79 | 119 | 36.8 | 45 | 13.9 | 74 | 22.9 | 34 | 10.5 | 85 | 26.3 |
| > 80 | 33 | 10.2 | 26 | 8.0 | 7 | 2.2 | 15 | 4.6 | 18 | 5.6 |
| **Marital status** |  |  |  |  |  |  |  |  |  |  |
| Single | 150 | 46.4 | 58 | 18.0 | 92 | 28.5 | 42 | 13.0 | 108 | 33.4 |
| Married | 173 | 53.6 | 46 | 14.2 | 127 | 39.3 | 39 | 12.1 | 134 | 41.5 |
| **Education (years)** |  |  |  |  |  |  |  |  |  |  |
| None | 23 | 7.1 | 16 | 5.0 | 7 | 2.2 | 8 | 2.5 | 15 | 4.6 |
| 1 - 6 | 137 | 42.4 | 56 | 17.3 | 81 | 25.1 | 44 | 13.6 | 93 | 28.8 |
| 7 and more | 163 | 50.5 | 32 | 9.9 | 131 | 40.6 | 29 | 9.0 | 134 | 41.5 |
| **Paid employment** |  |  |  |  |  |  |  |  |  |  |
| No | 231 | 71.5 | 77 | 23.8 | 154 | 47.7 | 60 | 18.6 | 171 | 52.9 |
| Yes | 92 | 28.5 | 27 | 8.4 | 65 | 20.1 | 21 | 6.5 | 71 | 22.0 |
| **Live alone** |  |  |  |  |  |  |  |  |  |  |
| No | 288 | 89.2 | 97 | 30.0 | 191 | 59.1 | 73 | 22.6 | 215 | 66.6 |
| Yes | 35 | 10.8 | 7 | 2.2 | 28 | 8.7 | 8 | 2.5 | 27 | 8.4 |
| **Smoking** |  |  |  |  |  |  |  |  |  |  |
| No | 300 | 92.9 | 97 | 30.0 | 203 | 62.8 | 71 | 22.0 | 229 | 70.9 |
| Yes | 23 | 7.1 | 7 | 2.2 | 16 | 5.0 | 10 | 3.1 | 13 | 4.0 |
| **Alcohol consumption** |  |  |  |  |  |  |  |  |  |  |
| No | 262 | 81.1 | 88 | 27.2 | 174 | 53.9 | 67 | 20.7 | 195 | 60.4 |
| Yes | 61 | 18.9 | 16 | 5.0 | 45 | 13.9 | 14 | 4.3 | 47 | 14.6 |
| **BMI (Kg/m<sup>2</sup>)** |  |  |  |  |  |  |  |  |  |  |
| Underweight ($\leq 19$) | 32 | 9.9 | 10 | 3.1 | 22 | 6.8 | 9 | 2.8 | 23 | 7.1 |
| Normal weight (20-24.9) | 201 | 62.2 | 58 | 18.0 | 143 | 44.3 | 46 | 14.2 | 155 | 48.0 |
| Overweight/Obesity ($\geq 25$) | 90 | 27.9 | 36 | 11.1 | 54 | 16.7 | 26 | 8.0 | 64 | 19.8 |
| **Comorbidity** |  |  |  |  |  |  |  |  |  |  |
| None | 178 | 55.1 | 53 | 16.4 | 125 | 38.7 | 46 | 14.2 | 132 | 40.9 |
| 1 | 115 | 35.6 | 30 | 9.3 | 51 | 15.8 | 21 | 6.5 | 60 | 18.6 |
| ≥2 | 30 | 9.3 | 21 | 6.5 | 43 | 13.3 | 14 | 4.3 | 50 | 15.5 |
| **Cognitive decline** |  |  |  |  |  |  |  |  |  |  |
| No | 172 | 53.3 | 43 | 13.3 | 129 | 39.9 | 45 | 13.9 | 127 | 39.3 |
| Yes | 151 | 46.7 | 61 | 18.9 | 90 | 27.9 | 36 | 11.1 | 115 | 35.6 |
| **Depression** |  |  |  |  |  |  |  |  |  |  |
| No | 199 | 61.6 | 50 | 15.5 | 149 | 46.1 | 43 | 13.3 | 156 | 48.3 |
| Yes | 124 | 38.4 | 54 | 16.7 | 70 | 21.7 | 38 | 11.8 | 86 | 26.6 |
| **Polypharmacy** |  |  |  |  |  |  |  |  |  |  |
| No | 159 | 49.2 | 36 | 11.1 | 123 | 38.1 | 39 | 12.1 | 120 | 37.2 |
| Yes | 164 | 50.8 | 68 | 21.1 | 96 | 29.7 | 42 | 13.0 | 122 | 37.8 |

SPPB, Short Physical Performance Battery; L-SPPB, low SPPB (score ≤7); H-SPPB, high SPPB (score >7); TL, telomere length; S-TL, short TL (≤4.22 kb pairs); L-TL, long TL (>4.22 kb pairs)

After the physical performance assessment, 104 participants (32.2%) were grouped in the low physical function (L-SPPB; score ≤7) group and 219 (67.8%) in the high physical performance (H-SPPB; score >7) group. Following the telomere length measurement, the short telomere length group (S-TL; ≤4.22 kb) included 81 participants (25.1%), while the long telomere length group (L-TL; >4.22 kb) 242 (74.9%) (Table 3).

**Table 3.**
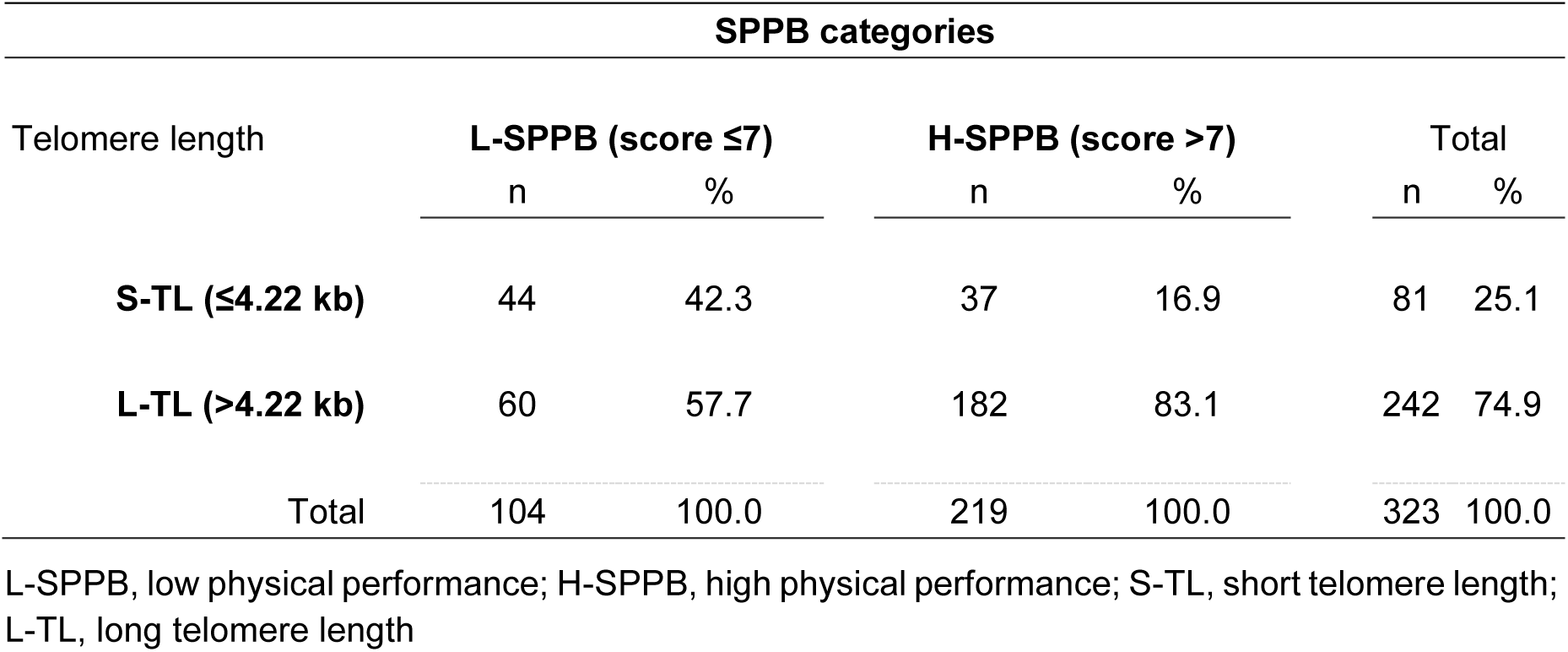
Physical performance categories by short and long telomere length.

We assessed the association between physical performance and telomere length and observed that participants in the low physical performance category (L-SPPB, score ≤7) had significantly shorter telomeres (_4.1_4.4_4.7_ mean and _3.5_4.0_4.5_ adjusted mean kb, p<0.001) while older adults in the high physical performance category (H-SPPBB, score >7) had longer telomeres (_5.5_5.7_5.9_ mean and _4.7_5.3_5.8_ adjusted mean kb, p<0.001) (Table 4), with a medium-to-high telomere length effect size (d= 0.762), according to an independent samples t-test: t(321)= 6.40, p<0.001; d= _0.521_0.762_1.002_; Cohen’s d for unequal group sizes with 95% CI).

**Table 4.** Association between physical performance and telomere length (n=323)

| Physical Performance | Total |  | Telomere length (kb pairs) |  |  |  |  |  |  |  |
| --- | --- | --- | --- | --- | --- | --- | --- | --- | --- | --- |
|  | n | (%) | LCI | Mean (95% CI) | UCI | p-value | LCI | adjMean (95% CI) | UCI | p-value |
| L-SPPB | 104 | 32.2 | 4.1 | 4.4 | 4.7 | <0.001 | 3.5 | 4.0 | 4.5 | <0.001 |
| H-SPPB | 219 | 67.8 | 5.5 | 5.7 | 5.9 |  | 4.7 | 5.3 | 5.8 |  |

L-SPPB, low SPPB score $\leq 7$ ; H-SPPB, high SPPB score $> 7$ ; LCI, lower confidence interval; UCI, upper confidence interval

Finally, our results indicate that the odds of being classified in the low physical activity category increase by _2.1_3.6_6.1_ times per kb of telomere attrition (adjOR _1.7_3.3_6.3_, p<0.001) compared to the high physical activity group (p<0.001) (Table 5).

**Table 5.** Association between physical activity and telomere length.

| Physical<br>Performance | Telomere length<br>(kb pairs) |  | OR<br>(CI 95%) | p-value | adjOR<br>(CI 95%) | p-value |
| --- | --- | --- | --- | --- | --- | --- |
|  | ≤ 4.22<br>% (n) | > 4.22<br>% (n) |  |  |  |  |
| SPPB score ≤7 | 13.6 (44) | 18.6 (60) | 3.6 (2.1-6.1) | <.001 | 3.3 (1.7-6.3) | <.001 |
| SPPB score >7 | 11.5 (37) | 56.3 (182) | 1<br>(Ref) |  |  |  |

SPPB, Short Physical Performance Battery; adjOR, adjusted OR

## Discussion

In this study, we examined the relationship between physical performance and telomere length in Mexican older adults under the hypothesis that participants with low physical performance would experience an accelerated telomere length attrition —indirectly observed as shorter telomere length—, compared to those with high physical function. Our results showed that older adults with low physical performance, considering standing balance, walking speed, and five times sit-to-stand (5xSTS) tests from the SPPB instrument, have shorter telomere length, while the opposite was observed in those participants with high physical function (Figure 2; [62]). This statistically significant difference between both groups corresponded to 1Kb (Table 5).

**Figure 2.**
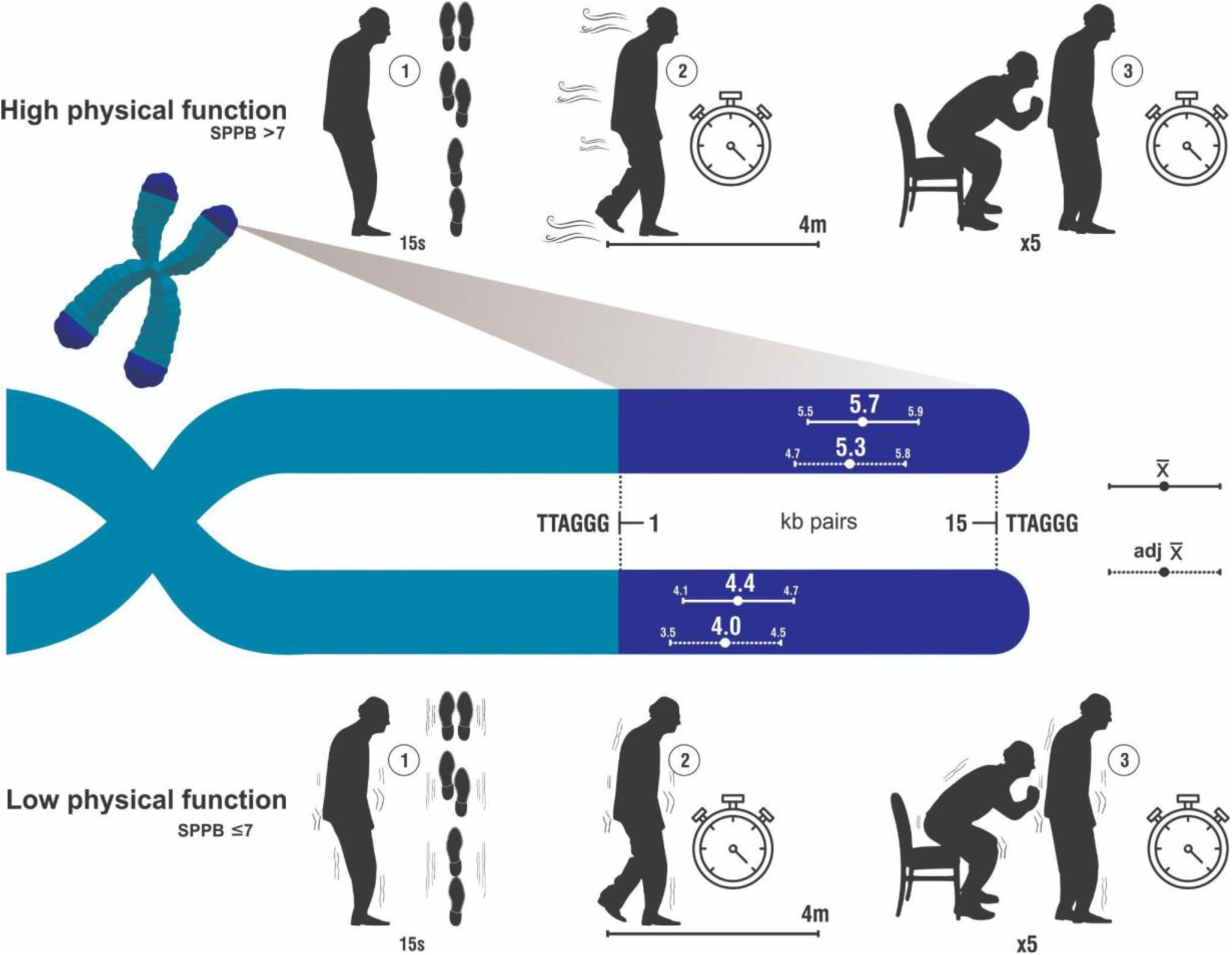
Relationship between physical performance and telomere length in older adults. This figure shows the positive relationship between physical function and telomere length [62].

Leukocyte telomere length (LTL) is a potential biomarker for aging [63] and may indicate the cumulative impact throughout life of environmental, lifestyle, and genetic factors; in fact, association studies have shown that longer telomeres predict a significant survival advantage in older adults [64]. Whether telomere length has a causal effect on mortality or is only a symptom of aging remains to be determined [65]. Previous potential mechanisms through which physical activity or exercise could affect telomere length have been proposed: telomerase activity, oxidative stress, inflammation, and skeletal muscle satellite cell content [44].

The combination of the degree and rate of change of each physiological and molecular mechanism is likely unique for different individuals, resulting in a heterogeneous aging phenotype [66] and varying susceptibility to develop multiple chronic conditions in old age.

The association between physical activity-inactivity and telomere length has been explored in healthy individuals [48,67], young professionals, and middle-aged athletes from track and field disciplines [68], and individuals with clinical conditions [69]. Moreover, physical inactivity is associated with short telomere length in normal conditions [48], as well as clinical conditions such as coronary heart disease [69], while moderate physical activity [67] and superior fitness levels observed in athletes [68] may provide a protective effect.

Any extreme activity is associated with shorter telomeres, even exercise. In addition, chronic stress [28], lack of physical activity [29], nutrition [30], smoking [31], pollution components of ambient air [32], and organophosphate insecticides [34] have been factors associated with shorter telomeres. All these factors can contribute, in turn, to a diminished physical capacity. On the other hand, physical activity maintains telomere length [70,71], probably by increasing telomerase reverse transcriptase expression and telomerase activity, thereby promoting telomere maintenance and preventing cellular senescence and aging phenotypes [72]. Also, physical activity is known to improve mental health and quality of life [73]. In adults, it has been reported that walking 2.5 hours per week is associated with longer telomeres compared to non-walkers and those who walk less time per week [74]. For either cause, we know that shorter telomeres could be a valuable prognostic biomarker of a worst outcome [75] that can be used to assess and monitor interventions designed to increase the individual’s physical activity.

A previous study used data from nine UK cohort studies to investigate telomere attrition between two measurements in older adults. While the authors interpreted their results as weak evidence of an association between leukocyte telomere length and physical performance [76], they did find significant differences between telomere length and physical performance at baseline. In contrast, a recent longitudinal 10-year follow-up study [77] examined the association between leisure-time physical activity (LTPA) and leukocyte telomere length (LTL) in 1,014 subjects with a mean age of 60.8 years at baseline. They found that, at baseline, the volume of LTPA was not associated with LTL. Nevertheless, the authors did find that attrition increased only in older women with high LTPA at baseline but not in men. In another study from Finland, the authors also observed shorter telomere length at follow-up and greater telomere attrition during follow-up time associated with poorer physical performance after adjusting for covariates (age at baseline, smoking status, body mass index at baseline, follow-up time, and educational attainment) in women only [9]. These findings suggest that the association between physical activity and telomere length may be sex-specific. However, as mentioned in our methods, we controlled for the potential confounding effect of sex; hence, our results indicate that the relationship between telomere length and physical performance is not merely due to differences in sex.

Moreover, a study on older Brazilians reported that subjects with lower telomere length had better physical capacity [53]. Discrepancies in telomere length and physical function measurements may arise from various factors, such as environmental influences and ethnicity, as recently reported for cardiovascular conditions [78] or even in sex-specific patterns [77]. However, in this Brazilian study, the participants were from the community, maintained good cognitive function with controlled comorbidities, and were actively involved in rural, domestic, and manual activities that require muscular, aerobic, and balance conditioning [53]. After adjusting for sociodemographic and health behavior covariates, a study conducted on a national representative sample of U.S. adults aged between 20 and 84 years old found that increasing one hour per week of vigorous leisure-time physical activity (LTPA) was associated with longer leukocyte telomere length (LTL), while the same increase in physical activity of household/yard work was associated with shorter LTL. However, no significant association was found for transportation physical activity or moderate LTPA with LTL [79].

One limitation of our study is the absence of previous physical activity and sedentary time to evaluate a longitudinal attrition rate due to this variable.

Considering these results, it is reasonable to hypothesize that a daily physical activity intervention, particularly vigorous leisure-time physical activity, could mitigate the factors that affect physical performance, which is also reflected in telomere length (TL) measurements. Hence, TL could be used as a biomarker to monitor and evaluate early interventions for a better functional aging process after adjusting for previous and current physical performance. These interventions should be prioritized for younger adults (between 50-60 yr) with shortened telomere length, regardless of the attrition-accelerating agent, to prevent or ameliorate its detrimental effects on physical activity.

## Conclusion

The study findings reveal a notable association between reduced physical function and shorter telomere length in Mexican older adults. It is crucial to acknowledge that various factors, including environmental influences, ethnicity, and sex, can influence comparisons of telomere length with physical activity. Given these potential disparities, there is a compelling imperative for future research, including longitudinal experiments to assess telomere length attrition both with and without physical activity while considering the diverse forms of physical activity in daily life. This research will yield valuable insights into the use of telomere length as a biomarker for monitoring and evaluating early interventions aimed at enhancing the process of healthy aging.

## Acknowledgments

The authors thank Aurelio García Cortés for his help in producing the figures.

This article is part of the requirements for JDME to obtain a Ph.D. in Biological Sciences at the Posgrado en Ciencias Biológicas, Universidad Nacional Autónoma de México (UNAM).

We thank the Consejo Nacional de Humanidades, Ciencia y Tecnología (CONAHCYT) of Mexico, for the scholarship awarded to JDME (CVU/Becario: 412887/263531).

## Contributorship

Conceptualization: JDME, PG and SSG; Data curation: JDME, MOR, and VGC; Formal analysis: JDME; Investigation: JDME, MOR, and SSG; Methodology: JDME, MOR, and SSG; Project administration: PG and SS; Resources: PG and SSG; Software: JDME; Supervision: JDME, PG and SSG; Validation: JDME, VGC and SSG; Visualization: JDME; Writing – original draft: JDME and MOR; Writing - review & editing: JDME, MOR, PG, VGC and SSG;

All authors approved the final version of the article.

